# Meta-analysis of diurnal transcriptomics reveals strong patterns of concordance and discordance in mouse liver

**DOI:** 10.1101/2022.06.22.497209

**Authors:** Thomas G. Brooks, Aditi Manjrekar, Gregory R. Grant

## Abstract

The accumulation of public transcriptomic timeseries data enables robust meta-analyses that were not possible until recently. To assess the consistency of biological rhythms across studies, 43 public mouse liver tissue timeseries totaling 805 RNA-seq samples were obtained and analyzed. Only the control groups of each study were included, in order to create comparable data. Technical factors in RNA-seq library preparation were the largest contributors to transcriptome-level differences, beyond biological or experiment-specific factors such as lighting conditions. Core clock genes were remarkably consistent in phase across all studies, while phase distributions of other periodic genes were generally less consistent. Overlap of genes identified as rhythmic across studies was generally low, up to around 50% between some of the highest sample count studies. Distributions of phases of significant genes were remarkably inconsistent across studies, but genes consistently identified as rhythmic clustered near ZT0 and ZT12 in acrophase. Data was integrated across studies in a JIVE analysis, which showed that the top two components of joint within-study variation are determined by time of day. A shape-invariant model with random effects was fit to the genes to identify the underlying shape of the rhythms, consistent across all studies. This revealed the extent of asymmetric and multimodal genes.

## Introduction

Circadian or diurnal transcriptomic experiments study changes in expression of the entire transcriptome as a function of the time of day. Individual studies are limited by the difficulty and expense of gathering a sufficiently large number of samples to power the required statistical analysis. However, a growing number of such data is now available in public repositories. While an increasing number of transcriptomics meta-analyses are being performed^1,2^, metanalyses examining the diurnal rhythm component of the transcriptome have so far been limited in scope^3–5^ and have utilized primarily older microarray experiments^6^. Here we expand these investigations to RNA-seq with particular attention to identifying and quantifying consistency of rhythmic behavior across studies.

While many available timeseries transcriptomics datasets investigate specific conditions and therefore contain non-comparable data, most include ‘control’ conditions that are nominally identical. By allowing the inclusion of similar, but not identical, conditions (such as both nighttime-restricted feeding and ad libitum feeding in mice), a large set of ‘control’ timeseries can be assembled. We investigate mouse liver - the most common mammalian tissue for circadian transcriptomics - by analyzing 43 RNA-seq timeseries studies containing 805 samples identified from the Gene Expression Omnibus (GEO) repository. For each study we started from the raw sequencing data which we processed in a uniform manner to obtain comparable quantifications from each timeseries. We assessed the consistency of these profiles using JTK_CYCLE analyses of the rhythmicity in each study.

We also performed additional meta-analyses across all studies. Meta-analyses often rely on random effects to capture differences across studies^7^. While most often used in the context of repeated-measures, there are random effects models, called shape-invariant models^8^ which perform non-linear curve fitting. We employ these models with random effects accounting for differences between timeseries from different studies to assess the shapes of the rhythmic profiles of genes that are consistent across studies.

## Results

We searched the GEO repository to identify mouse liver RNA-seq studies referencing the following terms: ZT, CT, zeitgeber, Bmal1, Cry1/2, Per1/2, Dbp, clock, constant conditions, entrain, darkness, circadian, or rhythm. This returned 275 GEO records, of which 33 have evenly spaced time points in mouse liver with resolution of at least 6 hours and span at least one full day. After selecting for approximate ‘control’ conditions and separating out by experiment, condition, and sex, we obtained 43 studies containing 805 samples, see Table 1 and Methods. Starting from sequencing reads, we quantified all samples using Salmon^9^, giving all data a consistent reference genome and annotation.

**Table 1.** Studies Analyzed. Studies were included if they had in-vivo mouse liver RNA-seq, had at least 4 evenly spaced timepoints per day, included data from ‘control’ conditions (meaning no major interventions or non-wild type genotypes, see methods), and the data was available on GEO and was compatible with our pipeline. Replicates column denotes the mean number of biological replicates at each collection time. Cycles denotes the number of 24-hour periods measured, counted such that, for example, samples every four hours for six timepoints would be one cycle (even though the ZT24 time would not be included until the seventh timepoint). Sequencing Type column indicates details the type of RNA-seq performed. M = male; F = female; LD = 12-12 hour light:dark conditions; DD = constant darkness conditions; GSE = Gene Expression Omnibus series identifier; SS = strand-specific; US = unstranded; SE = single-end; PE = paired-end; 3prime = 3-prime specific sequencing; PolyA = poly(A) selected sequencing; RiboZero = RiboZero rRNA depleted sequencing

| Name | GSE | PUBMED | Sex | Light | Age (weeks) | Sample Count | Timepoints Per Cycle | Replicates | Cycles | Sequencing Type | Notes |
| --- | --- | --- | --- | --- | --- | --- | --- | --- | --- | --- | --- |
| Benegiamo18<sup>20</sup> | GSE98042 | 29358041 | M | LD | 14 | 12 | 12 | 1 | 1 | SS SE RiboZero |  |
| Brooks22<sup>21</sup> | GSE115264 | 34724846 | M | DD | 16 - 24 | 25 | 6 | 4 - 5 | 1 | SS PE PolyA | Bmal1<sup>fx/fx</sup> |
| Cajan16 | GSE61775 |  | M | LD | 10 - 12 | 24 | 12 | 2 | 1 | US PE PolyA |  |
| Chaix19A<sup>22</sup> | GSE102072 | 30174302 | M | LD | 24 | 12 | 6 | 2 | 1 | SS SE PolyA | high fat diet |
| Chaix19B<sup>22</sup> | GSE102072 | 30174302 | M | LD | 24 | 11 | 6 | 1 - 2 | 1 | SS SE PolyA | night-restricted feeding; high fat diet |
| Du14<sup>23</sup> | GSE57313 | 24867642 | M | LD | 12 - 24 | 12 | 6 | 2 | 1 | US SE RiboZero | Alb<sup>Cre-ERT2</sup>; tamoxifen treated |
| Fader19<sup>24</sup> | GSE119780 | 31015483 | M | LD | 9 - 10 | 24 | 8 | 3 | 1 | SS SE PolyA | sesame oil gavage |
| Gaucher19<sup>25</sup> | GSE132103 | 31757851 | M | LD | 9 - 24 | 18 | 6 | 3 | 1 | SS SE PolyA |  |
| Greenwell19A<sup>26</sup> | GSE118967 | 30995463 | M | LD | 12 - 13 | 18 | 6 | 3 | 1 | SS SE 3prime |  |
| Greenwell19B<sup>26</sup> | GSE118967 | 30995463 | M | LD | 12 - 13 | 18 | 6 | 3 | 1 | SS SE 3prime | night-restricted feeding |
| Guan20<sup>27</sup> | GSE143524 | 32732282 | M | LD | 8-12 | 24 | 8 | 3 | 1 | SS PE PolyA | Rev-erbα<sup>fl/fl</sup>; Rev-erbβ<sup>fl/fl</sup> |
| Hirako18<sup>28</sup> | GSE109908 | 29805094 | F | LD | 10 | 12 | 4 | 3 | 1 | SS SE PolyA |  |
| Kinouchi18<sup>29</sup> | GSE107787 | 30566858 | M | LD | 8 | 18 | 6 | 3 | 1 | SS SE PolyA |  |
| Lahens15<sup>30</sup> | GSE40190 |  | M | DD | 6 | 8 | 4 | 1 | 2 | US PE PolyA |  |
| Levine20<sup>31</sup> | GSE133989 | 32369735 | M | LD | 32 | 19 | 6 | 3 - 4 | 1 | SS SE PolyA |  |
| Li19A | GSE113745 |  | M | LD | 16 | 12 | 6 | 2 | 1 | SS SE PolyA |  |
| Li19B | GSE113745 |  | M | LD | 76 | 12 | 6 | 2 | 1 | SS SE PolyA |  |
| Li20<sup>32</sup> | GSE133342 | 32160860 | M | DD | 6 | 6 | 6 | 1 | 1 | US PE RiboZero | six weeks of DD |
| Manella21<sup>33</sup> | GSE159135 | 34059820 | M | LD | 12 - 16 | 24 | 12 | 2 | 1 | SS SE 3prime | Alb-Cre<sup>+</sup> |
| Mermert18<sup>34</sup> | GSE101423 | 29572261 | M | LD | 8 - 12 | 6 | 6 | 1 | 1 | SS SE PolyA |  |
| Mortimer21<sup>35</sup> | GSE117134 | 34036284 | F | LD | 8 - 12 | 18 | 6 | 3 | 1 | SS PE RiboZero | Alfp-Cre<sup>-/-</sup>/tg |
| Morton20 | GSE151565 |  | M | LD | 26 | 77 | 8 | 5 - 6 | 1.625 | SS PE PolyA |  |
| Pan20<sup>36</sup> | GSE130890 | 31935211 | M | DD | 8 - 12 | 48 | 12 | 2 | 2 | SS PE PolyA | XBP1<sup>fllox</sup> |
| Quagliarini19A<sup>37</sup> | GSE108688 | 31706703 | M | LD | 17 - 18 | 17 | 6 | 2 - 3 | 1 | US PE PolyA |  |
| Quagliarini19B<sup>37</sup> | GSE108688 | 31706703 | M | LD | 17 - 18 | 18 | 6 | 3 | 1 | US PE PolyA | high fat diet |
| Quagliarini19C<sup>37</sup> | GSE108688 | 31706703 | M | LD | 17 - 18 | 17 | 6 | 2 - 3 | 1 | SS PE PolyA |  |
| Quagliarini19D<sup>37</sup> | GSE108688 | 31706703 | M | LD | 17 - 18 | 17 | 6 | 2 - 3 | 1 | SS PE PolyA | high fat diet |
| Rubio-Ponce21<sup>38</sup> | GSE125867 | 33937766 | M | LD | 8 - 9 | 18 | 6 | 3 | 1 | US SE PolyA |  |
| Sinturel17A<sup>39</sup> | GSE73552 | 28475894 | M | LD | 12 - 14 | 48 | 12 | 4 | 1 | SS PE RiboZero |  |
| Sinturel17B<sup>39</sup> | GSE73552 | 28475894 | M | LD | 12 - 14 | 24 | 6 | 4 | 1 | SS PE RiboZero | night-restricted feeding |
| Stubblefield18<sup>40</sup> | GSE105413 | 29386110 | M | LD | 9 - 12 | 32 | 8 | 4 | 1 | SS SE RiboZero |  |
| Trott18<sup>41</sup> | GSE36871 | 29300726 | M | LD | 12 - 24 | 12 | 6 | 2 | 1 | US SE PolyA |  |
| Weger19A<sup>42</sup> | GSE114400 | 30344015 | M | LD | 15 - 16 | 12 | 6 | 2 | 1 | SS PE RiboZero |  |
| Weger19B<sup>42</sup> | GSE114400 | 30344015 | F | LD | 15 - 16 | 12 | 6 | 2 | 1 | SS PE RiboZero |  |
| Weger21A<sup>43</sup> | GSE135898 | 33452134 | M | LD | 9 - 14 | 12 | 6 | 2 | 1 | SS PE PolyA |  |
| Weger21B<sup>43</sup> | GSE135875 | 33452134 | M | LD | 9 - 14 | 12 | 6 | 2 | 1 | SS PE PolyA |  |
| Weger21C<sup>43</sup> | GSE135898 | 33452134 | M | LD | 9 - 14 | 12 | 6 | 2 | 1 | SS PE PolyA |  |
| Wu19<sup>44</sup> | GSE138019 | 31875550 | M | LD | 16 - 24 | 6 | 6 | 1 | 1 | SS SE RiboZero |  |
| Xin21<sup>45</sup> | GSE150380 | 33889826 | F | LD | 9 | 28 | 6 | 4 | 1.17 | SS PE RiboZero | night-restricted feeding |
| Yang16A<sup>46</sup> | GSE70497 | 26843191 | M | DD | 16 - 24 | 24 | 6 | 4 | 1 | US PE PolyA | Bmal1<sup>fl/fl</sup>; tamoxifen treated |
| Yang16B<sup>46</sup> | GSE70499 | 26843191 | M | LD | 6 - 14 | 9 | 6 | 1 - 2 | 1 | SS PE PolyA |  |
| Yang16C<sup>46</sup> | GSE70499 | 26843191 | F | LD | 6 - 14 | 9 | 6 | 1 - 2 | 1 | SS PE PolyA |  |
| Zhang14<sup>47</sup> | GSE54651 | 25349387 | M | DD | 6 - 7 | 8 | 4 | 1 | 2 | SS PE PolyA |  |

To better assess differences between studies, we note that the most common study design is male mice, LD lighting, and ad libitum feeding of standard chow without any interventions. There are 20 such studies which are therefore highly comparable, differing primarily in age or factors that are often unreported (such as housing).

### Technical factors dominate biology and study design

To assess the overall similarities of studies, we performed a PCA on the log-scaled transcripts per million (log TPM) expression values. These revealed that the largest differences between studies were driven by technical factors in the sequencing (Figure S 1). In particular, the three timeseries which sequence only the 3’ end of the transcript, were outliers. Similarly, RiboZero versus PolyA-selected libraries are clearly distinct. Smaller differences were observed between stranded versus unstranded sequencing and paired-end versus single-end sequencing. Differences from biological factors, such as male and female, were smaller than theses technical differences.

### Phases of core clock genes are consistent

Using the reported ZT/CT times for each study, we plotted time-course profiles of the TPM expression values of 7 core clock genes (Arntl, Cry1, Cry2, Per1, Per2, Nr1d1, and Clock)^10^ for each study and found that the peaks and troughs for these genes are well aligned across all timeseries (Figure 1, Figure S 2). While the amplitudes vary moderately, the plots reveal remarkable consistency in phase and period across all studies despite the differences in study designs.

**Figure 1.**
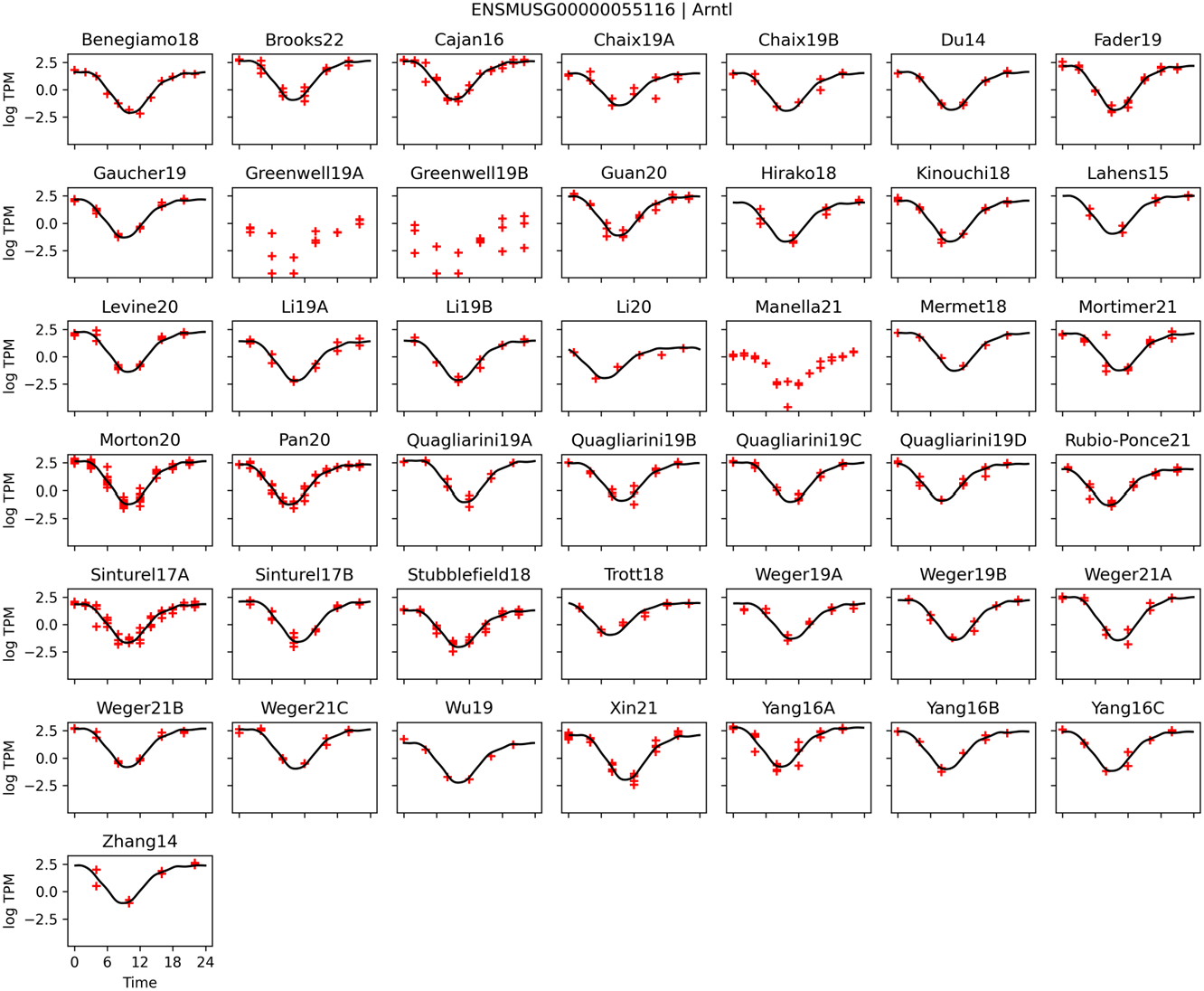
Core clock genes are highly consistent. Data from 40 mouse liver RNA-seq circadian timeseries were processed. Arntl (Bmal1) gene quantified transcripts per million (TPM) in red. Shape invariant models were fit to each gene with random effects allowing variation of amplitude, mesor, and phase between studies; study-specific fit line (black). Time 0 corresponds to lights on (ZT0 or CT0 depending on study) Shape invariant models curve fit in black (with three studies excluded from the fits due to using an uncommon sequencing methodology). See other core clock genes in Figure S 2.

### Single-study analyses have low consistency

We ran JTK_CYCLE^11^ (JTK) on each timeseries separately. We then compared the results of the datasets in several ways. First, we selected the top 1000 most rhythmic genes identified by JTK *p*-values in each timeseries and compared the overlaps of these lists between different studies (Figure S 3). Notably, no two studies have overlaps reaching 600 genes. Studies with large sample size (*n* > 26) typically have pairwise overlaps of 400-555 genes between each other and overlaps of 200-400 genes when compared to smaller studies. Small sample size studies (n < 18) have a modest agreement between each other of just 100-300 genes. The three studies with the largest overlaps of 533-555 are the three largest studies (Morton20, Pan19, and Sinturel17A) even though Pan19 differs in lighting conditions (12-hour light-dark and constant darkness). This emphasizes the importance of having enough samples to obtain repeatable results. Studies from the same laboratories showed only modestly higher agreement with each other than with other similar studies.

We next compared all studies with the single largest study, Morton20. In all studies, the fraction of genes identified as significant at FDR < 0.05 that were also significant in Morton20 ranged from 82% to 100% (median 94%). This indicates that most genes identified as significantly rhythmic in a study will reproduce in a sufficiently high-powered replication study.

We next compared consistency of the distribution of phases of identified rhythmic genes. These distributions are routinely used to summarize overall activity in the transcriptome. However, these showed remarkable inconsistency across studies (Figure 2). While many studies showed preferential clustering of phases near particular times of day, the locations of these clusters differed substantially between studies. These differences are not explained by factors such as 12-hour light-dark cycles (LD) versus constant darkness (DD) conditions; for example, Morton20 and Sinturel17A have the same conditions, large sample sizes, and high overlap in identified rhythmic genes, but have almost opposite peaks in their phase distributions. However, when a single set of rhythmic genes is chosen for all studies, then the phase distributions are considerably more consistent, with all showing a peak near ZT12 (Figure S 4). In contrast to phase, amplitude distributions were comparable across studies (Figure S 5).

**Figure 2.**
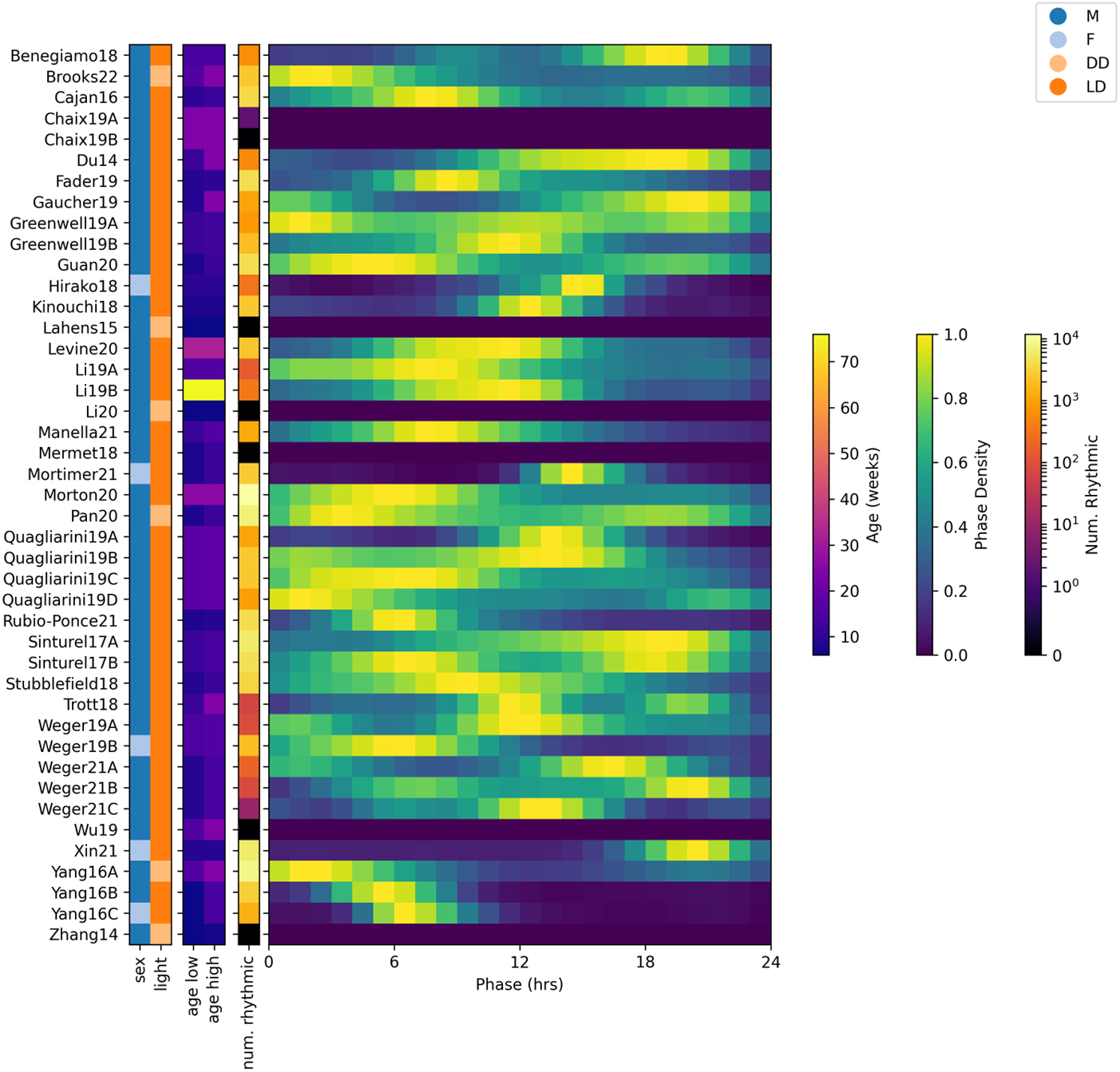
Consistency of phase distributions. JTK_CYLCE was run on each timeseries, and results were compared. Distributions of phases among genes identified rhythmic by JTK (at a Benjamini-Hochberg Q-value < 0.1), showing notable inconsistency between studies. Distributions are normalized to peak one, with the total number of genes identified as rhythmic shown in separate column. Phase distributions skipped in studies with fewer than 10 rhythmic genes. M=male; F=female; DD=constant darkness; LD=light-dark conditions.

To identify the genes consistently identified as rhythmic across studies, we employed a simple voting-counting metric. We defined the *robustness score* of a gene as the number of timeseries in which it was identified to be significantly rhythmic according to JTK (Figure 3 a), out of a max possible score of 43. We identified 4267 genes with a robustness score of at least 12, which has a *p* < 0.05 chance of giving even one false positive gene, see Methods. Moreover, 319 genes reached a robustness score of 30 or more and therefore exhibit rhythmic behavior across many studies, see Supplemental Table S1. High robustness genes cluster around ZT0 and ZT12 in acrophase (Figure 3 b).

**Figure 3.**
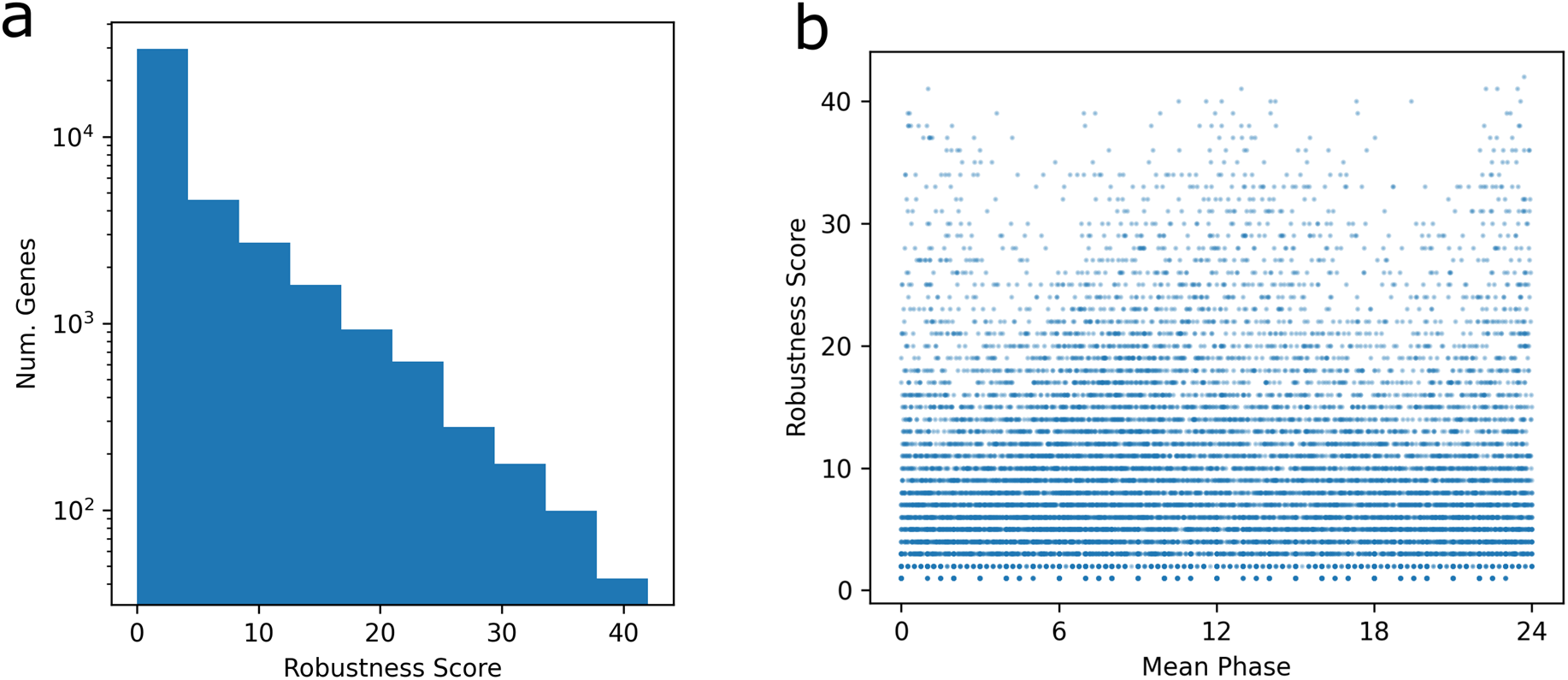
Robustness of genes. JTK_CYCLE results were summarized across all studies to identify genes that were highly consistent. The robustness score was computed as the number of studies in which the genes had JTK_CYCLE *p*-value under 0.05. Correcting for the number of studies and genes, robustness scores of 12 or higher have *p* < 0.05 if the gene is not rhythmic in any study. (a) Number of genes by robustness score. (b) Plot of robustness score by mean (across studies in which JTK_CYCLE *p* < 0.05) phase for each gene. Genes with the highest robustness scores cluster in phase near ZT0/ZT24 or ZT12.

### Shape-invariant models reveal the shape of the transcriptome rhythmicity

Shape-invariant models (SIMs) fits a flexible curve to multiple independently measured studies, assuming there is a consistent shape across studies^8^. Moreover, they allow for random-effects modelling by allowing each measured study to have differences in phase, amplitude, and mid-level (mesor). Applying this to our dataset allows the pooling of information from all studies to determine more accurately the underlying consistent shape of the rhythms within each study. We identified 2545 significantly rhythmic shapes (see Methods). Core clock genes were well-identified (Figure 1a, Figure S 2).

To visualize the overall spectrum of diurnal profiles, we normalized all SIM fits to have amplitude 1 and to peak at the same time. Then a t-SNE dimension-reduction was performed on these normalized shapes and the results were plotted (Figure 5a). Genes with significant fits were classified to either as monomodal with symmetric (n=1322) or asymmetric (n=1149) peaks, or as multimodal (n=74). This emphasizes the diversity of profiles present in the transcriptome, though the most common shapes are similar to the classic cosine curve.

**Figure 4.**
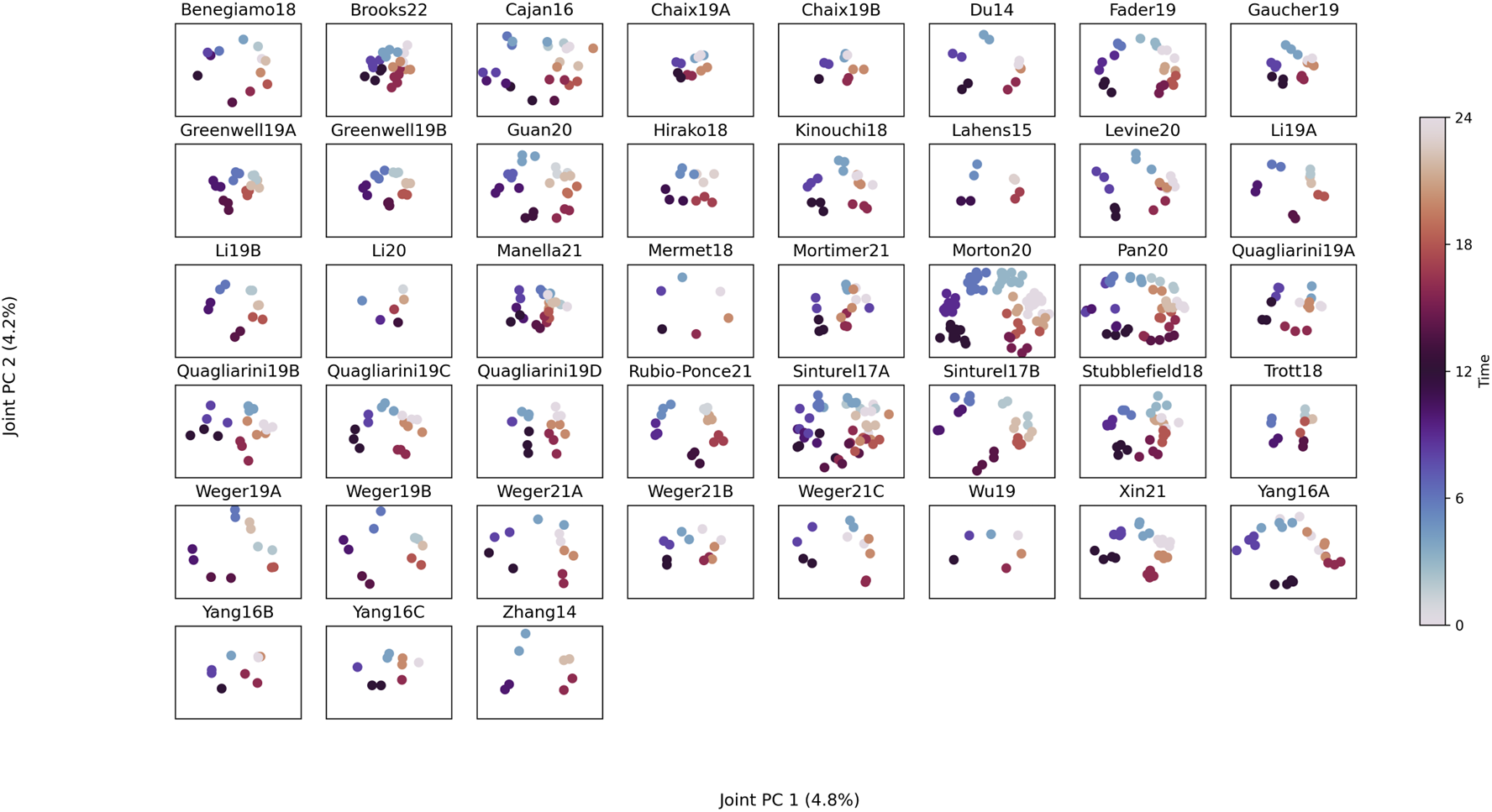
Consensus PCA reveals consistent rhythmicity across studies. A joint and individual variation estimation (JIVE) analysis was used to determined loadings of genes that have consistently high variance within each study (regardless of between-study differences). The identified two joint variance components are plotted in each timeseries, showing consistent separation by time-of-day. All plots use the same gene loadings.

**Figure 5.**
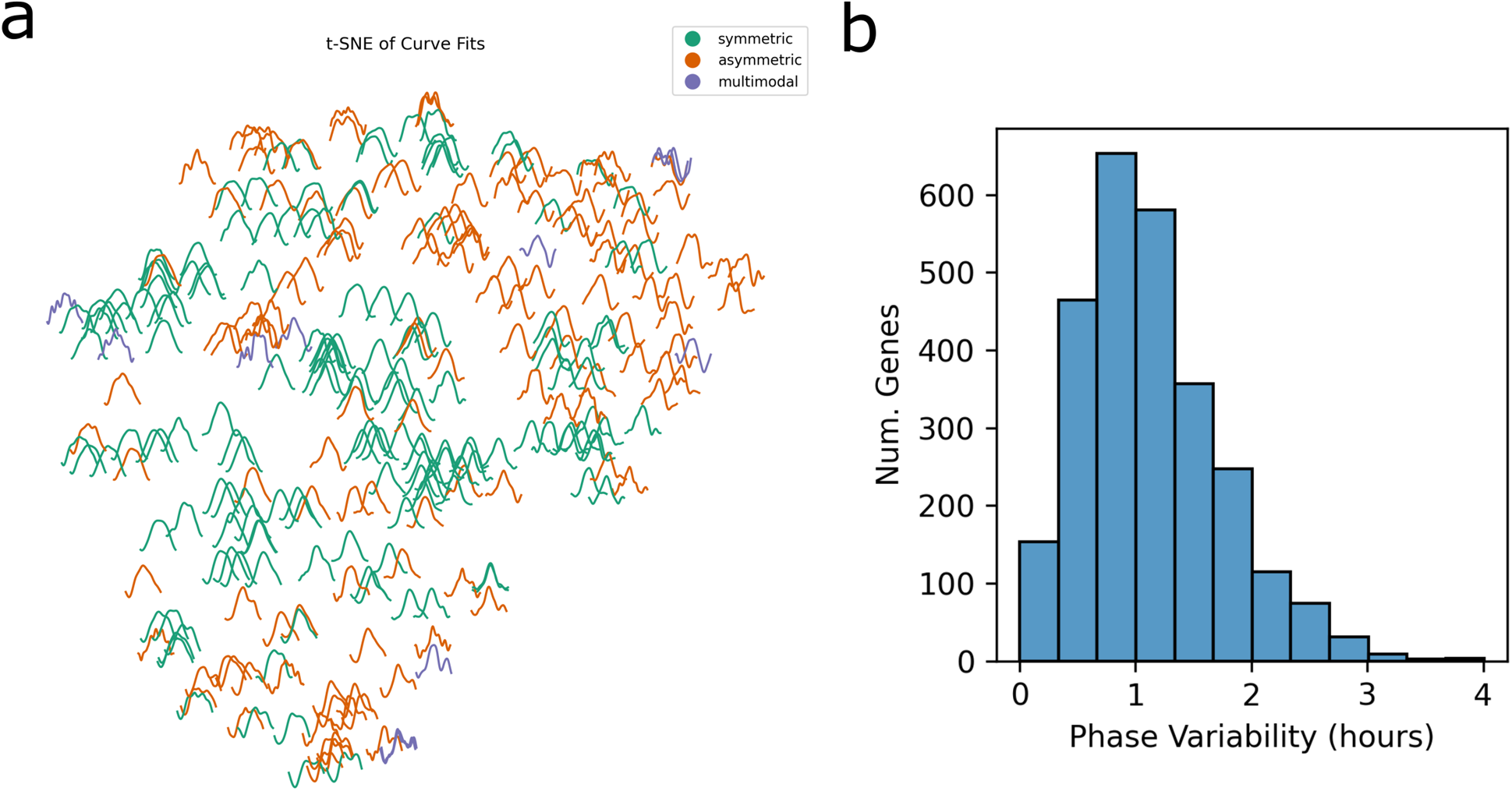
Shape Invariant Models. Shape invariant models were fit to each gene with random effects allowing variation of amplitude, mesor, and phase between studies. (a) t-SNE of fit curves of identified rhythmic genes (300 randomly sampled genes shown). Fit curves were normalized to have the same peak, mean, and amplitude. Genes were classified as symmetric (n=1322 genes), asymmetric (n=1149), or multimodal (n=74) according to the fit curve. (b) Each study is fit a unique phase variable, pooling appropriately across studies. Histogram of the variability of the phases in the rhythmic genes, measured as the fit phi SD value, converted to hours by the inverse logistic function.

### Joint PCA analysis (JIVE) identifies consistent rhythmicity across studies

To examine consistent factors of variance across the studies, a joint and individual variance estimate (JIVE) analysis was performed^12^. While PCA on aggregated data from multiple studies identifies variance primarily between studies (Figure S 1a), JIVE divides the within studies variance into a joint component (common to all studies) and individual components (distinct for each study). Specifying two components of joint variation and one per study of individual variation, the joint variance components capture the rhythmicity in factors that cleanly and consistently separate timepoints across all studies, see Figure 4. These two joint variance components together account for 9.0% of the within-study variance. In contrast, PCA run on individual studies gives inconsistent results, see Figure S 6.

### Genes with high variability in phase

Next, we considered the set of rhythmic genes which displayed the largest variability in their phases between studies. We hypothesized that these genes will be sensitive to external factors. Using the identified rhythmic SIM fits, the mean variability in phase was 1.1 hours, see Figure 5 b, determined by transforming the SIM phi value to hours by an inverse logistic function to approximate the standard deviation of phases. We identified 239 genes with at least 2 hours phase variability. Pathway analysis identified no pathways enriched for high phase variability.

### Consistent non-rhythmic genes

We searched for genes that were consistent across studies and time by requiring that for each gene, the mean TPM was at least 1, the standard deviation was at most half the mean TPM, and that JTK_CYCLE *q*-value was at least 0.05, as well as not having a rhythmic SIM fit. We chose these criteria to require high expression, low variance, and no detectable time-of-day dependence. A total of 108 genes met these criteria, see Supplemental Table S2. These genes represent candidate lists of ‘housekeeping’ genes that repeatedly have minimal time varying across a wide selection of studies and have high expression values. The commonly used housekeeping gene Gapdh was not on the list, due to having standard deviation of TPM over half of its mean TPM in three studies, as well as being significantly rhythmic by JTK in one study (and close to significant in several others). This demonstrates that these criteria are stringent and identify only highly consistent genes in mouse liver samples and may be useful as reference non-cyclic genes.

## Discussion

Meta-analyses of transcriptomics are becoming increasingly popular as more datasets become publicly available, and the importance and practicality of such studies are increasing. Here, we perform the first large-scale meta-analysis of circadian or diurnal timeseries transcriptomics that we are aware of. We found that analyses restricted to single studies consistently capture rhythms in core clock genes. However, substantial variability outside of the core clock between studies highlights the limitations of single-study datasets. Variability was such that differences from light condition (LD versus DD) were smaller than between-study variation, likely due to low sample counts. Despite this, we conclude that even low sample count studies primarily identify rhythmic genes that do reproduce in other, larger sample sizes studies.

Phase distributions showed marked differences between studies, even those with the largest sample counts or highest temporal resolutions and under the same conditions. This suggests caution while interpreting phase distribution plots from individual studies. Since phase distributions were considerably more consistent when a fixed set of genes was compared across all studies, the differences in phase distributions may be driven by differences in the set of genes identified as rhythmic rather than the phases of individual genes.

In contrast, meta-analysis identifies rhythmic factors that are consistent across many studies. A JIVE analysis demonstrates that times accounts for 9% of the within-study variance by identifying two components of variation common to all studies. These components give highly consistent results in all studies, demonstrating that despite their differences in single-study analyses, all studies contain a large underlying component of consistency.

Individual studies have limitations in resolution and replication necessary to confidently identify the shape of time-course expression profiles at the transcriptome-wide scale. By pooling data from all studies in a SIM analysis, we obtained reliable curve fits that do not overfit to the noise in any individual study. These allow us to observe that approximately half of all rhythmic genes in mouse liver have asymmetric patterns and a small number show multimodal patterns, even under light-dark conditions. Since many analysis methods make the assumption of symmetry, such as JTK_CYCLE^11^ and cosinor^13^, this informs the choice of alternative methods that do not require symmetry, such as RAIN^14^.

We observed large differences from sequencing parameters (such as rRNA depletion method or strand specificness), which were not always well-described in the corresponding publications. We therefore recommend more prominently describing such parameters in future studies.

One limitation of this study is the inclusion of data under multiple biological conditions. This likely decreases the amount of observed consistency between studies. However, even restricting to a subset of studies with highly consistent designs (male mice, LD lighting, and ad libitum feeding of standard chow) we find substantial inconsistencies on single-study analyses. Moreover, by including multiple biological conditions, results from SIM and JIVE analyses will capture effects that are consistent across those conditions and therefore of broad interest. Since these meta-analytic methods naturally account for differences between the individual studies, the inclusion of multiple study designs and conditions should not compromise the results.

## Conclusions

When a gene is found to be rhythmic in one data set and not in another, it can be due to technical factors that influence the statistical power to detect, or it can reflect the true biology – the gene in fact was rhythmic in one set of animals and not in the other. Datasets which compare one condition to a control condition wish to identify effects that are driven by the condition. However, the results of this meta-analysis indicate that a significant amount of variation can be due to variation in the baseline themselves. This then raises the question of the proper interpretation of differences identified between condition and control experiments.

We have observed from this study that rhythm detection in transcriptomics is limited by sample counts, by demonstrating considerable variability in ‘control’ conditions across studies in the identified rhythmic genes as well as in distributions of phase among those genes. Meta-analysis is a key tool for researchers to arrive at consensus that remains under used in circadian transcriptomics despite a growing wealth of data available. Meta-analysis also answers key questions that are not answerable in any individual study no matter the sample count, such as: how similar would our observations be if we someone else repeated the experiment? Effects that exist in a single study may be of limited interest, even those that pass statistical significance. Use of ‘mega-analysis’^15^, where the original data from all studies is analyzed instead of just summary statistics, further allows exploring new questions, such as robustly identifying the shape of the temporal profile of these genes.

## Methods

### Data collection

We searched the GEO repository using the GEOmetadb R package^16^ version 1.44.0 to identify mouse liver RNA-seq data containing references to the following terms: ZT, CT, zeitgeber, Bmal1, Cry1/2, Per1/2, Dbp, clock, constant conditions, entrain, darkness, circadian, or rhythm. The search was performed on GEO data collected 2021-07-09. The resulting accessions were assessed for the following: timeseries data of mouse liver samples; evenly spaced timepoints; at least one cycle of data; resolution of at least 6 hours or faster; data were compatible with our pipeline (excludes color-space data or datasets with large adapter sequences that did not align well); containing at least one ‘control’ condition. Control conditions were defined as meeting the following:

- Genotype: either wild-type mice (usually C57BL/6J) or a genotype used as control to another genotype (for example, a Cre+ genotype)
- Feeding: either ad libitum food and water or night-restricted (ZT12-ZT14) feeding
- Sex: any allowed; if both male and female, sexes were separated into separate timeseries
- Light conditions: either light-dark (12hr:12hr) or constant darkness conditions
- Interventions: none during days of sample collection; placebo or control treatments completed at least 24-hour before sample collection

Some GEO records contain multiple matching experiments, such as two control timeseries for comparisons with two different genotypes. In such cases, the individual timeseries were treated as separate studies. Studies were labelled according to the first author and year of publication, or according to author and year made public on GEO if no publication was yet available.

### RNA-Seq data processing

A snakemake (v6.5.3) pipeline was developed to reproducibly process the data^17^. Reads were downloaded as `sra` files and converted to `fastq` format using the `efetch, prefetch`, and `fastq-dump` commands from `edirect` (v15.3) and `sratoolkit` (v2.11.0). Starting from sequencing reads, we quantified all samples using Salmon^9^ (v1.4.0) to the GRCm38.75 Mus musculus transcriptome with the `-k 31` index option. Salmon was run with the `-l A --softclip --softclipOverhangs --seqBias --gcBias --reduceGCMemory --biasSpeedSamp 10 --posBias -p` options as well as `-g` to quantify at the gene-level.

This generated read count estimates and transcripts per million (TPM) counts for each of 40614 genes or transcripts.

### Quality Control

All data was manually inspected for consistent read depth and alignment statistics within each timeseries. Outlier data were identified by performing principal component analysis on each timeseries and removing any samples that were at least three standard deviations from the mean in the first principal component. If any outliers were discarded, this process was repeated on the remaining samples until no more outliers were detected. In total, 6 samples were identified as outliers and removed from further analysis.

### Rhythmicity testing

JTK_CYCLE^11^ was run on each timeseries TPM data with 24 hour periods using MetaCycle 1.2.0^18^. Benjamini-Hochberg *q*-values were computed from JTK_CYCLE *p*-values after first dropping any genes that had mean read depth less than 2 across all samples in the timeseries. Dropped genes were assigned *q*=1.

Since the default for JTK_CYCLE is to use 20-28 hour periods, we also ran it with that setting to determine if non-24 hour genes were detectable. We selected the 6 studies which have 8 or more timepoints per day (to improve ability to identify period) and are the most under the most common conditions (male, LD, ad libitum feeding). For each pair of studies, we computed Cramer’s V and Spearman R statistics comparing JTK periods on the genes significant in both (q < 0.05). These showed low consistency, with medians of V = 0.10, R = 0.07 and no pair achieving higher than V=0.19 or R=0.18. To check for consistently low-period genes, we searched for genes with period less than 24 in at least four of these six studies and with no period 24 or greater (when *q* < 0.05). Only 13 genes were identified. However, 100 random permutations of the period values gave a median of 16 identified genes by the same criteria. Due to this observed inconsistency in non-24 hour period estimates, reported results were exclusively from using the fixed 24 hour period, which also matches best-practice recommendations^19^.

### Robustness score

For each gene, we computed its robustness score as the number of timeseries in which it was identified as significantly rhythmic according to JTK_CYCLE *p* < 0.05. With 43 total timeseries, the average nonrhythmic gene is expected to have a robustness score of about 2. With 40,614 genes measured and conservatively assuming that all were non-rhythmic, we can calculate that the Bonferroni-corrected *p*-value of getting even a single gene with robustness score at least 12 is 0.035, from the survival function of the binomial distribution with n=43, p=0.05. Therefore, all genes with at least robustness score 12 are assumed to be genuinely rhythmic (in at least some studies) at a family-wise error rate of less than 0.05.

### PCA analyses

All PCA analyses were run on TPM data transformed by *log*(*x*+0.01). PCA was run on the joined data from all timeseries to assess differences between studies. Furthermore, we ran PCA on each individual study to assess PCA performance as it would happen in any individual study, see Figure S 6. Finally, we ran a JIVE^12^ analysis, grouping within individual studies. We used two components of joint variation (as that typically is needed to capture the circular effect of time-of-day) and one component of individual variation in the JIVE analysis, to allow between study variations.

### Shape-invariant models

Shape-invariant models (SIM) were run to fit timeseries data to a periodic spline with a random-effects model allowing each timeseries in the meta-analysis to have different amplitudes, phases and mean values (mesors). The random effects structure compensates for the expectation that different studies have different values while still prioritizing consistency between studies. Fit splines then identify the consistent shape of curves throughout the day across all studies. Periods were fixed to 24 hours, though the spline may also fit ultradian rhythms at harmonic periods (such as 12 or 8 hour periods). The R package `assist` (v3.1.7) was used with the `snm` function^8^. The convergence condition was set to the “PRSS” method with convergence criterion `prec.out` = 0.05. Values for `lambda` parameter were restricted to 10^-4^ to 10^3^ to prevent overfitting that happened at the default settings (10^-10^ to 10^3^). The SIM model was fit to each gene that had non-zero values in at least 1/3^rd^ of all samples across all studies, after outlier removal described earlier.

The R^2^ goodness-of-fit value was computed for each timeseries and each gene as (SS_total_ – SS_resd_)/SS_total_ where SS_total_ and SS_resid_ and the squared sum of differences from the mean value and from the model fit of the data, respectively. Due to the existence of random effects in the model, this is not guaranteed to be positive, unlike the R^2^ of a linear model. Nonetheless, larger values are indicative of better fits.

The fit for a gene was classified as rhythmic if it satisfied:

- Fit converged in at most 30 iterations,
- SD of the logAmp random effect parameter is at most 3 (i.e., most timeseries should have amplitudes within a factor of 20 of the mean amplitude),
- `funcDf` < 15 (i.e., fitting at most 15 degrees of freedom to the spline, considering that 23 degrees of freedom is enough to fit each hour with an independent value),
- `median_t` > 2 (i.e., the median across time of the *t*-statistic (difference from zero divided by standard error) is at least 2, so that the spline is significantly non-zero at most times), and
- median across all studies of R^2^ > 0.25.

To estimate the false discovery rate of these criteria, a one-shot permutation test was performed, where the data within each study had timepoint labels permuted randomly. Permutations were done independently in each gene, but due to computational costs only a single permutation was taken for each gene. In the permuted data, there were only 15 genes satisfying the rhythmicity criteria while in the original data there were 2545 such genes. Therefore, we estimate an FDR of less than 0.01.

Fits were classified as either monomodal symmetric, monomodal asymmetric, or multimodal. Symmetry was assessed by comparing the fit curve to all its cyclic mirror images (one reflected about every quarter hour). If all mirror images contained some points of at least 2 standard errors (determined by the pstd variable) different from the original curve, then the gene was classified as asymmetric. Therefore, fits symmetric about even one point would not be identified. To identify multimodal genes, we looked for at least two distinct peaks, meaning two points at least two standard errors from 0 and with at least one point below zero between them in both clockwise and counterclockwise directions.

## Data Availability

All data from this study was obtained from GEO and a Snakemake pipeline to download and process these and create all figures in the study is available at https://github.com/tgbrooks/circadian_comparison. Since significant computational time goes into processing these datafiles, all quantified, labelled data, metadata and JTK and SIM results, are provided in Supplemental File 1, which would be sufficient for many future analyses.

## Acknowledgments

This research was supported by funding from the National Center for Advancing Translational Sciences Grant (5UL1TR000003) and was started as part of the American Physician Scientists Association’s Virtual Summer Research Program 2021.

TGB received funding from Calico Laboratories. The authors declare no other conflicts of interest.

## Supplemental Files

Table S1: Robustly Rhythmic Genes

List of genes with robustness core at least 30, indicating highly robust rhythms in most studies.

Table S2: Highly Consistent Non-Rhythmic Genes

List of highly consistent, non-rhythmic genes (gene symbol and Ensembl gene ID) with low sample variance, for potential use as reference genes or “true nulls” in rhythmic analyses.

File S1: Collected Data and Metadata

Compilation of all the quantified data, both as TPM and as counts, as well as sample metadata (study ID and time-of-day), study metadata (see Table 1) and results from JTK and SIM analyses. This data should be sufficient for future meta-analysis of these datasets.

