## Supplemental Figures for "Meta-analysis of diurnal transcriptomics reveals strong patterns of concordance and discordance in mouse liver"

#### Supplemental Figures

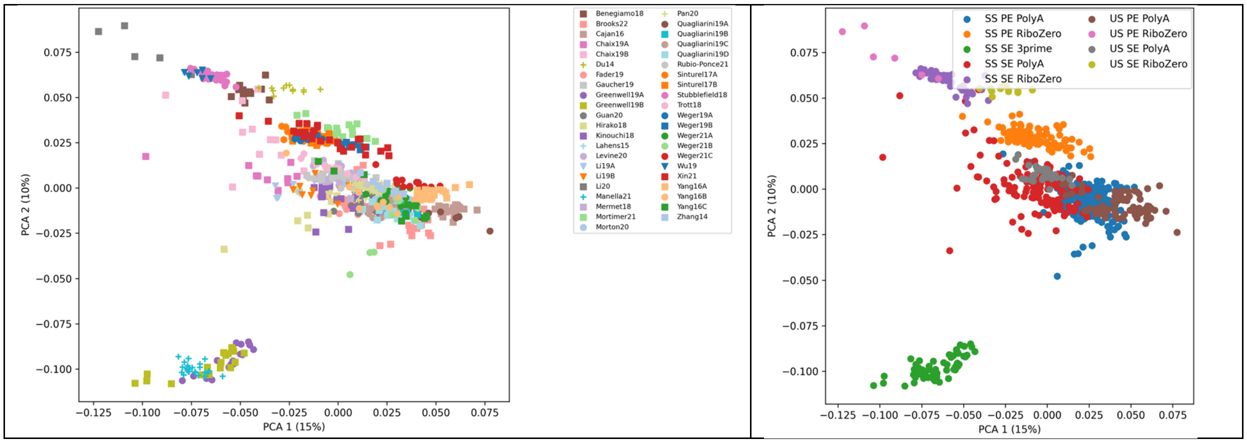


##### Figure S 1 PCA of study data

Principal component analysis of RNA-seq from 805 mouse liver samples from 43 different timeseries. (left) PCA labelled by the timeseries each sample came from. (right) Labelled by the sequencing type of the study, revealing that the largest differences are due to technical choices. SS=strand-specific; US=unstranded; PE=paired-end; SE=single-end

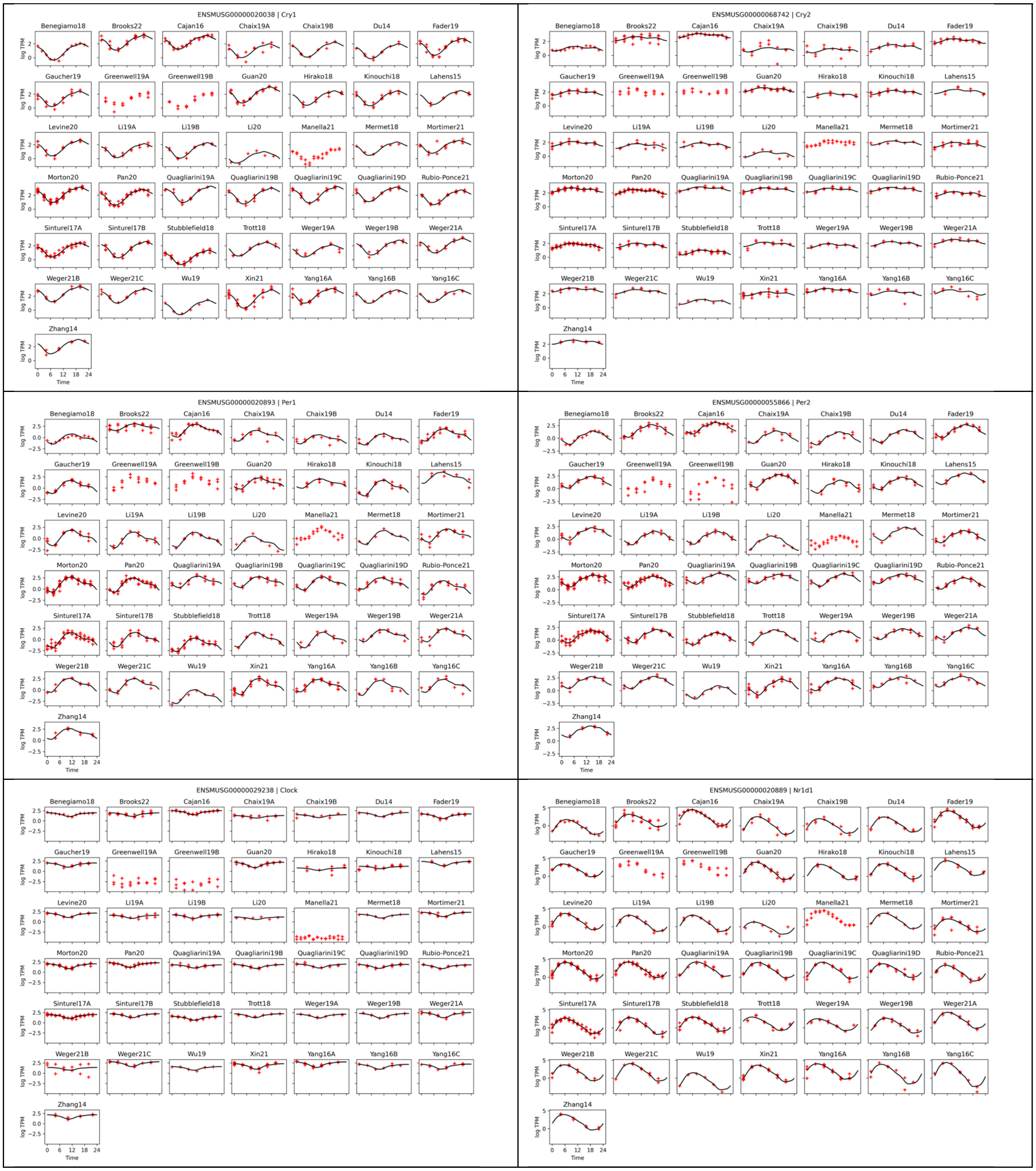


#### Figure S 2 Core clocks gene expression profiles

Six core clock gene (Cry1, Cry2, Per1, Per2, Clock, and Nr1d1 from top left to bottom right) expression levels by time, in each of the 43 studies. Red dots are log TPM levels and black curves are the SIM fits (with three studies excluded due to unusual sequencing methodology). All clock genes show consistent shapes and phases across studies.

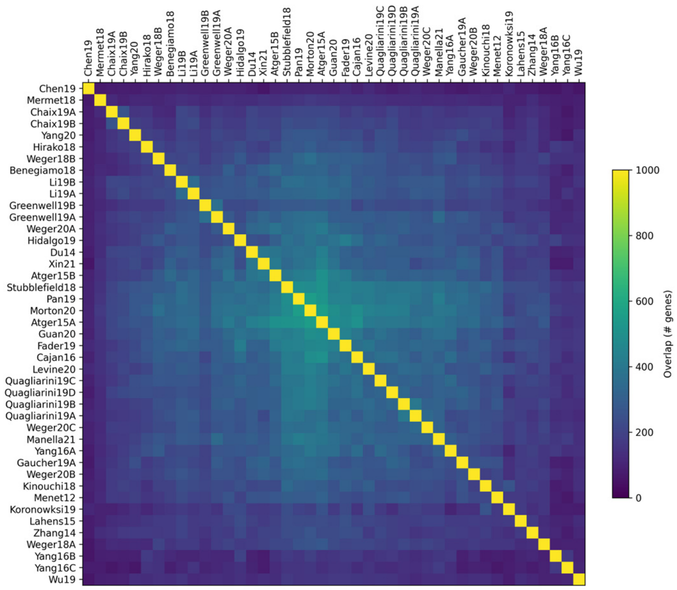

#### Figure S 3 JTK Rhythmic Genes Overlaps

JTK_CYLCE was run on each timeseries, and results were compared. Number of genes overlap between the top 1000 most rhythmic (by JTK *p*-value) in each study.

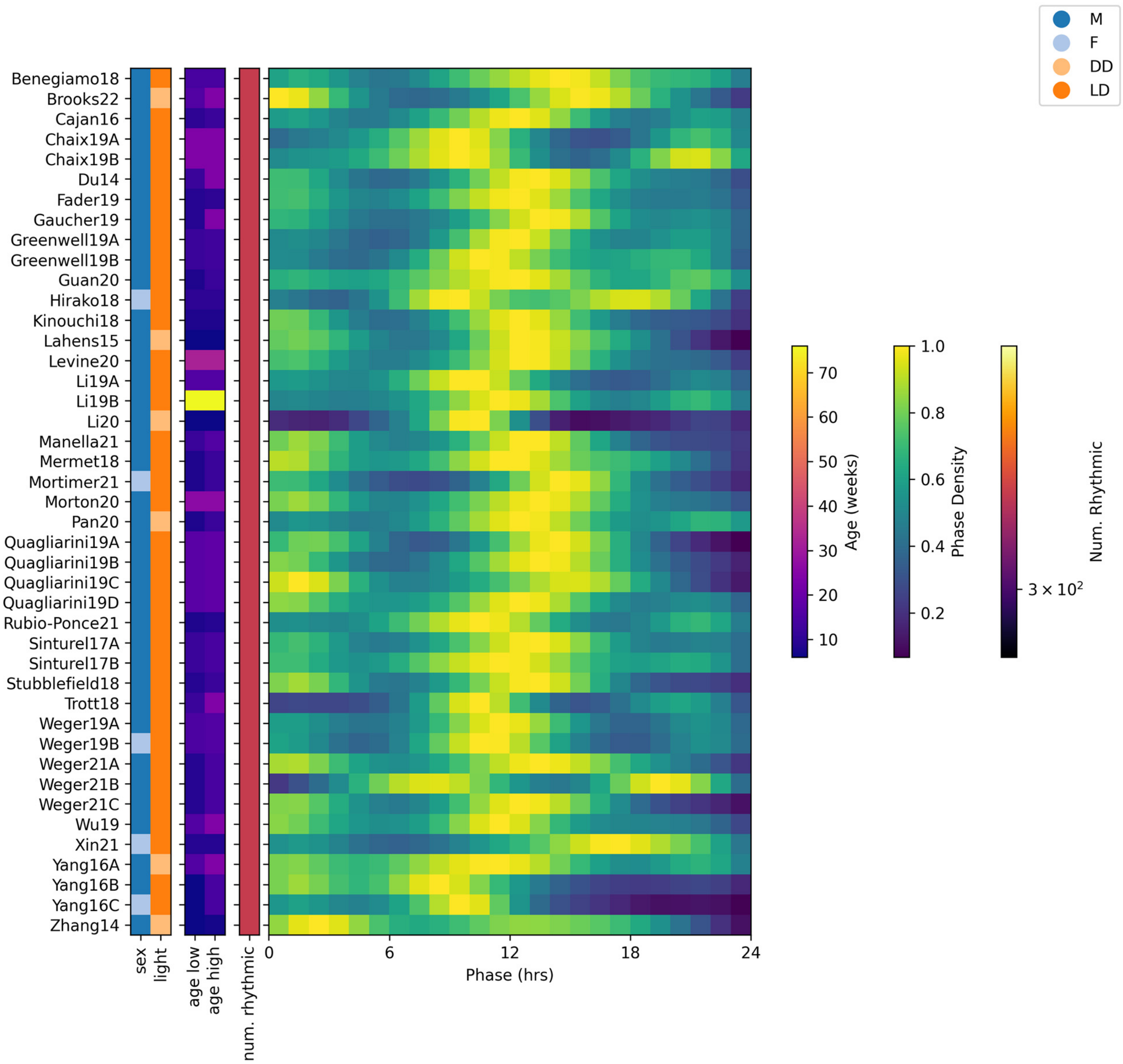


#### Figure S 4 JTK Phase distribution in select genes

JTK_CYLCE was run on each timeseries, and results were compared. Distributions of phases among a fixed set of 319 genes that were consistently rhythmic ($p < 0.05$ in at least 30 studies). Compared to Figure 2, these show considerably higher consistency across studies Distributions are normalized to peak one, with the total number of genes identified as rhythmic shown in separate column. Phase distributions skipped in studies with fewer than 10 rhythmic genes. M=male; F=female; DD=constant darkness; LD=light-dark conditions.

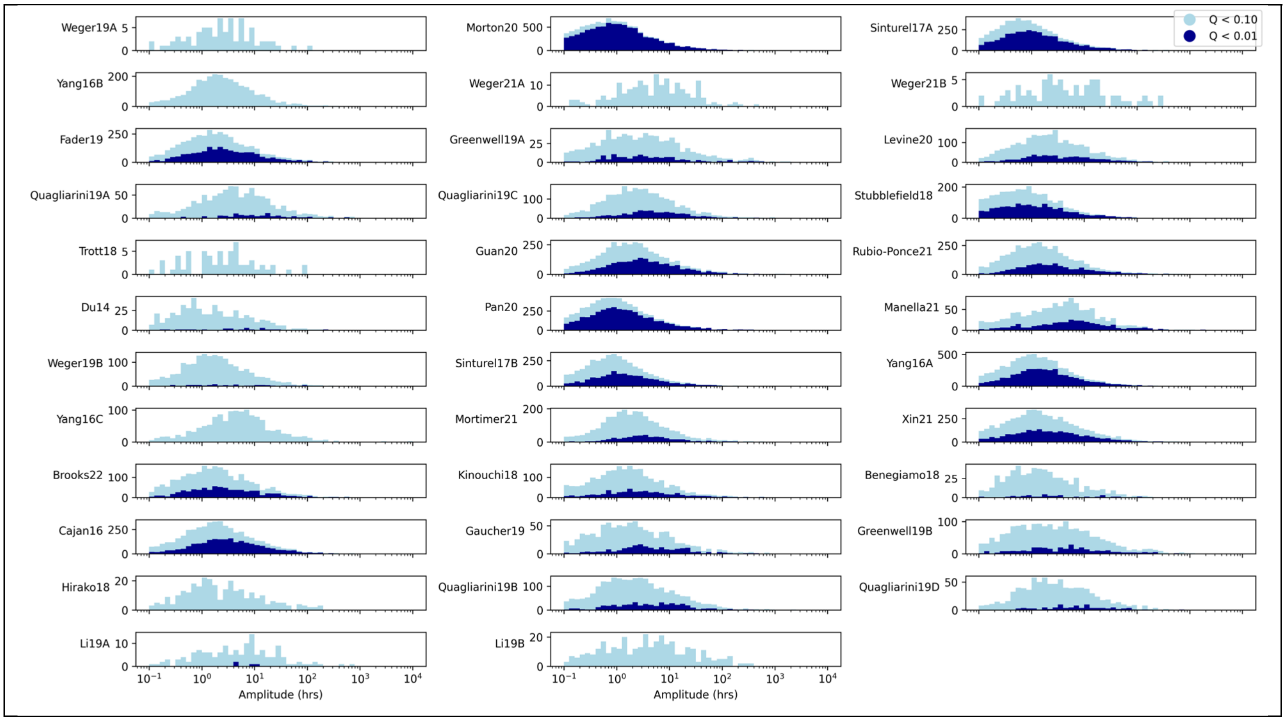


#### Figure S 5 JTK amplitude distributions

Amplitude distributions from JTK_CYCLE results, among significantly rhythmic genes at two significance cutoffs (by shade).

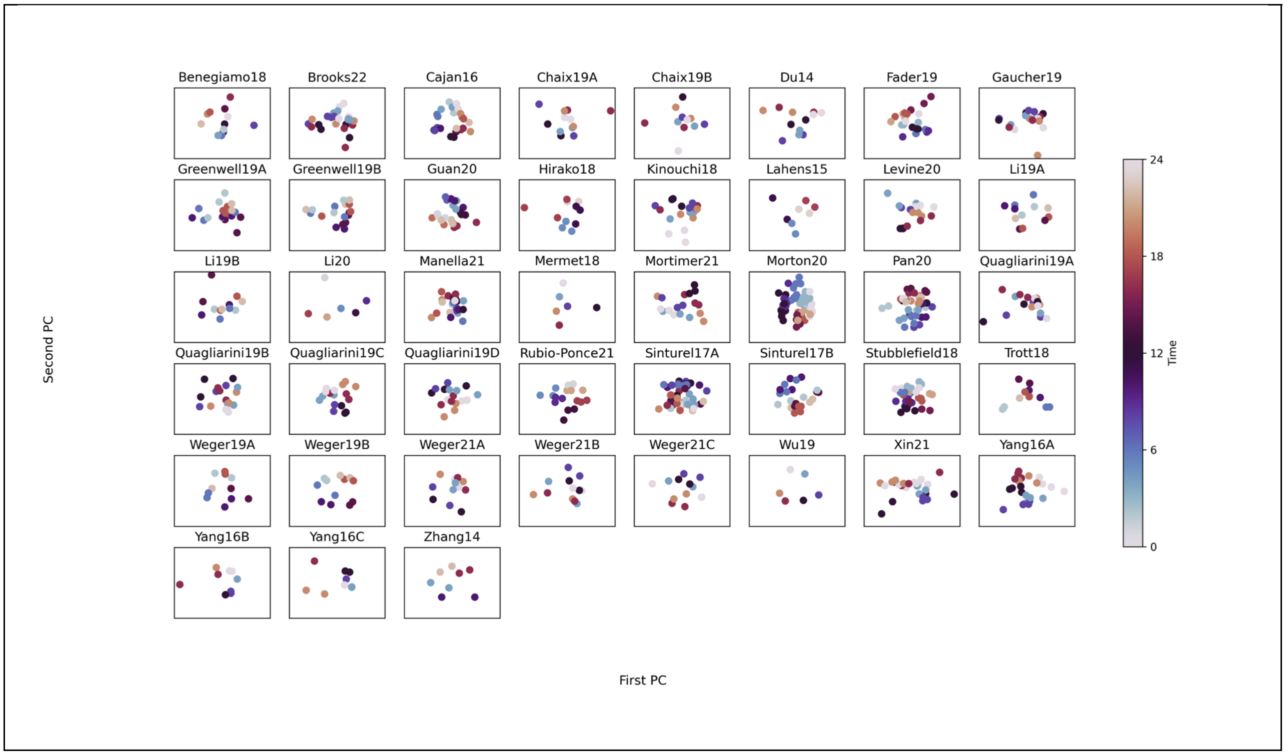


Figure S 6 Individual PCA

PCA was performed on each study individually and the top two components were plotted. Unlike JIVE, PCA components are unrelated in distinct studies, and these top two components do not reliably capture time variation within a study and are not comparable between different studies.

### Supplemental Files

#### Table S1: Robustly Rhythmic Genes

List of genes with robustness core at least 30, indicating highly robust rhythms in most studies.

#### Table S2: Highly Consistent Non-Rhythmic Genes

List of highly consistent, non-rhythmic genes (gene symbol and Ensembl gene ID) with low sample variance, for potential use as reference genes or “true nulls” in rhythmic analyses.

#### File S1: Collected Data and Metadata

Compilation of all the quantified data, both as TPM and as counts, as well as sample metadata (study ID and time-of-day), study metadata (see Table 1) and results from JTK and SIM analyses. This data should be sufficient for future meta-analysis of these datasets.
