## Supplemental File 1 for "Meta-analysis of diurnal transcriptomics reveals strong patterns of concordance and discordance in mouse liver": README.txt

"Meta-analysis of circadian transcriptomics reveals strong patterns of concordance and discordance in mouse liver"
Thomas G. Brooks, Aditi Mandjrekar, Gregory R. Grant

Supplemental File 1: DATA AND METADATA

CONTENTS:

DATA FILES:
    tpm.by_sample.txt.gz:
    num_reads.by_sample.txt.gz:

        These gzipped tab-separated tables contain quantified expression values for all genes quantified across all studies.
        No filter or normalization is performed, values are directly as reported by Salmon.
        Both TPM and NumReads values are displayed.

        "Name" column gives to the Ensembl gene or transcript identifier
        "Symbol" column gives the gene symbol (if any) for the gene
        All other columns give read counts of the sample with that ID
        Use sample_metadata.txt to map those to study and time-of-day values

    sample_metadata.txt:

        Tab-separated table containing metadata of every sample quantified

        "sample_id" columns gives the GEO identifier of the sample
        "study" gives the ID of the study (see study_metadata.txt)
        "time" column gives the ZT or CT time of the study, as the study reported it
                Note that this is not constrained to 24 hours and not all studies
                start at time 0
        "outlier" if True, then was identified as an outlier within the study

    study_metadata.txt:

        Tab-separated table of data about the studies, such as sequencing type and sex of mice

        "Name" columns gives the study ID (for joining on sample_metadata.txt)


RESULTS FILES:
    jtk.results.txt.gz:

        Gzipped tab-separated results table from running JTK with a fixed 24 hour period on each studies.

        "ID" gives Ensembl gene identifier
        "study" is the study ID
        "BH.Q" is JTK-reported Q value, run on all genes (see also 'qvalue' column)
        "ADJ.P" is JTK-reported p-value
        "PER" is JTK-reported period (always 24 if significant)
        "LAG" is JTK-reported phase
        "AMP" is JTK-reported amplitude
        "dropped" indicates if the gene had low expression in a study and was dropped from q-value calculations
        "qvalue" is the BH q-value for the genes in a study that were not dropped (or 1 if dropped)
                In contrast to 'BH.Q' which includes all genes (even non-expressed) and so has lower power

    robustness_score.txt:

        Generated gene robustness scores, i.e. the count of number of studies in which the JTK p-value was < 0.05
        Higher scores indicate consistently rhythmic genes.
        Values above 12 are statistically unlikely to occur by chance even in a single gene.
        Values much higher than 12 indicate that it must be rhythmic in many studies.

        'ID' columns gives gene Ensembl ID
        'score' gives the robustness score

SIM FITS:
    SIM/summary.txt:

        Main summary of gene-level data of SIM fits

        "gene" gives Ensembl gene identifier
        "fit_peak_time" gives time (0-24) of peak of the average fit
        "fit_trough_time" gives time (0-24) of trough of the average fit
        "fit_amplitude" gives amplitude (in log scale TPM) of the average fit
        "fit_mesor" gives the average mesor (mid point, in log scale TPM) of the average fit
        "mesor_stderr" gives standard error fo the mesor value
        "re_logAmp_sd" gives the random effects of the amplitude (in log-of-log scale TPM) standard deviation across studies
        "re_phi_sd" gives the random effects of phi (the logit-transformed phase) standard deviation across studies
        "re_mesor_sd" gives the random effects of mesor (in log scale TPM) standard deviation across studies
        "lambda" gives the smoothing parameter
        "sigma" gives the estimated residual size of the fit
        "iteration_times" gives number of iterations before convergence
        "iteration_converg" gives whether the iteration converged
        "funDf" gives number of degrees of freedom used in the function fit
        "log_likelihood" gives log likelihood of the fit
        "AIC" gives the AIC of the fit
        "BIC" gives BIC of the fit
        "t_stat" gives the maximum over the fit curve of the t statistic (difference from mesor over standard error)
        "median_t" gives median over the fit curve of the t statistic
        "rel_amp" gives relative amplitude (amplitude over mesor)
        "rsquared" gives R-squared of the fit
        "is_rhythmic" denotes if the gene was classified as significantly rhythmic
        "category" gives which category the gene was classified into based off shape
        "num_peaks" gives the number of identified peaks in the shape of the fit (if is_rhythmic)

    SIM/curves_fit.txt
        Shape of the fit curves.

        "gene" gives the gene Ensembl ID
        All other columns give the value of the fit at a specified time t from 0 to 1.
        These curves are relative to the mesor.
        To obtain the fit of a specific study, need to apply the following:

        u = ((t + (alogit(study_phi) - 0.5)) % 1)
        study_fit = exp(study_logAmp) * fit_value + mesor + study_mesor

        where study_phi, study_logAmp, study_mesor the random effect fits for that study from re.txt
        and t is the time being fit (column name of curves_fit.txt file) and fit is the entry in curves_fit.txt
        and alogit is the inverse logistic transform
        See scripts/plot_spline_fits.py in the github repo for an example use.

    SIM/curves_pstd.txt:
        Same as curves_fit.txt but instead of reporting the fit shape, reports the PSTD standard error estimate by time

    SIM/re.txt:
        Random effects fits of each study

        "gene" gives gene ID
        "study" gives study ID
        "logAmp" gives study logAmp fit (in log-of-log TPM scale)
        "phi" gives the logit-transformed phase
        "mesor" gives the mesor value (in log TPM scale)

    SIM/re_structure.txt:
        Gives the random effect structure, i.e. correlations between the three fitted random effects for each gene
